# *KairoSight-3.0*: A Validated Optical Mapping Software to Characterize Cardiac Electrophysiology, Excitation-Contraction Coupling, and Alternans

**DOI:** 10.1101/2023.05.01.538926

**Authors:** Kazi T Haq, Anysja Roberts, Fiona Berk, Samuel Allen, Luther M Swift, Nikki Gillum Posnack

**Affiliations:** Sheikh Zayed Institute for Pediatric Surgical Innovation, Children’s National Health System, Washington DC, USA 20010; Children’s National Heart Institute, Children’s National Hospital, Washington DC, USA 20010; Department of Biomedical Engineering, School of Engineering and Applied Sciences: George Washington University, Washington DC, USA 20037; Department of Pediatrics, Department of Pharmacology & Physiology, School of Medicine and Health Sciences: George Washington University, Washington DC, USA 20037

**Keywords:** Cardiac electrophysiology, calcium handling, excitation-contraction coupling, optical mapping

## Abstract

**Background:** Cardiac optical mapping is an imaging technique that measures fluorescent signals across a cardiac preparation. Dual optical mapping of voltage-sensitive and calcium-sensitive probes allow for simultaneous recordings of cardiac action potentials and intracellular calcium transients with high spatiotemporal resolution. The analysis of these complex optical datasets is both time intensive and technically challenging; as such, we have developed a software package for semi-automated image processing and analysis. Herein, we report an updated version of our software package (*KairoSight-3*.*0*) with features to enhance characterization of cardiac parameters using optical signals.

**Methods:** To test software validity and applicability, we used Langendorff-perfused heart preparations to record transmembrane voltage and intracellular calcium signals from the epicardial surface. Isolated hearts from guinea pigs and rats were loaded with a potentiometric dye (RH237) and/or calcium indicator dye (Rhod-2AM) and fluorescent signals were acquired. We used Python 3.8.5 programming language to develop the *KairoSight-3*.*0* software. Cardiac maps were validated with a user-specified manual mapping approach.

**Results:** Manual maps of action potential duration (30 or 80% repolarization), calcium transient duration (30 or 80% reuptake), action potential and calcium transient alternans were constituted to validate the accuracy of software-generated maps. Manual and software maps had high accuracy, with >97% of manual and software values falling within 10 ms of each other and >75% within 5 ms for action potential duration and calcium transient duration measurements (n=1000-2000 pixels). Further, our software package includes additional cardiac metric measurement tools to analyze signal-to-noise ratio, conduction velocity, action potential and calcium transient alternans, and action potential-calcium transient coupling time to produce physiologically meaningful optical maps.

**Conclusions:** *KairoSight-3*.*0* has enhanced capabilities to perform measurements of cardiac electrophysiology, calcium handling, and the excitation-contraction coupling with satisfactory accuracy.

**Graphical Abstract Demonstrating Experimental and Data Analysis Workflow:** Created with Biorender.com

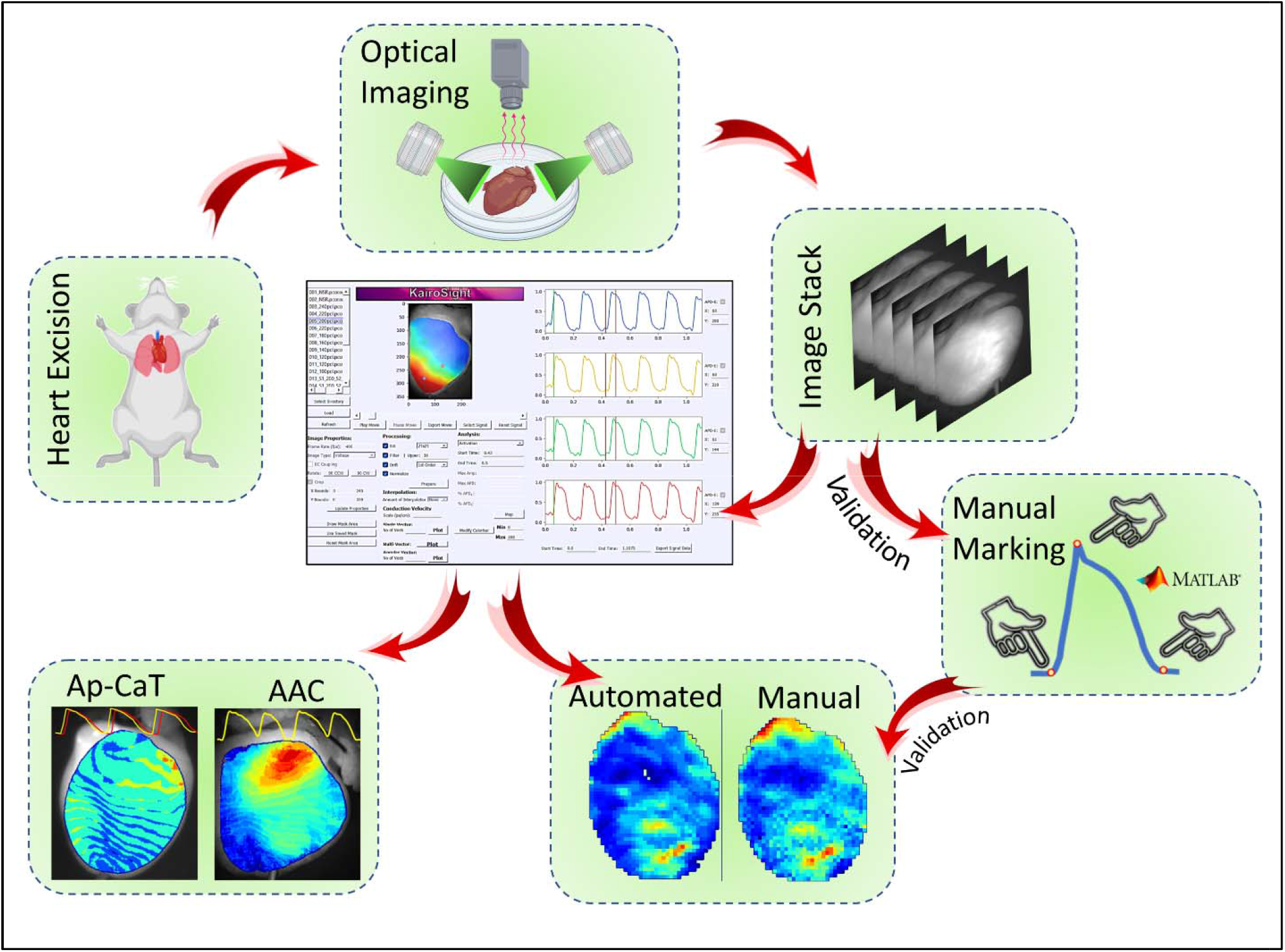

## 1. INTRODUCTION

Recent advances in cardiac electrophysiology research have been propelled by optical mapping, a powerful imaging technique that is used to monitor dynamic changes in metabolic status, transmembrane potential, and/or intracellular calcium concentration^1–3^. Most commonly, cardiac action potentials (AP) and calcium transients (CaT) are imaged simultaneously from cardiac tissue through the combined use of voltage-sensitive and calcium-sensitive fluorescent dyes or genetically-encoded indicators (with complementary spectral properties)^4–8^. To characterize the properties of these fluorescent signals, a scientific camera with high spatiotemporal resolution is used to capture electrical propagation and intracellular calcium cycling at a high sampling frequency. Since optical mapping studies require a fast rate of acquisition and fluorescent signals have a small fractional change, the acquired images often require post-processing to improve the signal-to-noise ratio (SNR) quality^5,9,10^. Accordingly, the analysis and interpretation of optical mapping data is both time- and resource-intensive, which has prompted the development of several unique software packages for semi-automated image processing and analysis (e.g., *Rhythm*^11^, *ElectroMap*^12^, *ORCA*^13^, *COSMAS*^14^, *Spiky*^15^). A common feature across these software packages is the ability to generate epicardial action potential (AP) activation and repolarization maps. Uniquely, *COSMAS*, and *ElectroMap* provide calcium transient duration, amplitude, and alternan maps; *RHYTHM* enables 3D data and arrhythmia analysis; *ORCA, COSMAS*, and *ElectroMap* provide conduction velocity measurements; *Spiky* allows for both whole heart and cardiomyocyte signal analysis. Since each software package has its own features, researchers may be required to utilize multiple approaches to comprehensively analyze an optical dataset.

We recently developed an open-source software package to analyze cardiac optical mapping data using a graphical user interface, which aimed to facilitate broader use by individuals with limited data analysis expertise. The described software was aptly named *KairoSight*^16^ in reference to the Greek word for “opportune time” (Kairos) and the ability to “see” voltage or calcium signals acquired from cardiac tissue. Based on feedback from the research community, we have expanded the available features in an updated software package entitled *KairoSight-3*.*0*. Herein, we validated the accuracy of our automated software using a user-directed algorithm developed in MATLAB (MathWorks, Natick MA). Further, we expanded the available analysis features to include: 1) a user-defined masking tool, 2) SNR mapping tool, 3) signal duration measurements using updated algorithms, 4) AP and CaT alternan maps, 5) AP-CaT coupling maps, 6) conduction velocity (CV) module, and 7) AP (AP duration) and CaT (CaT duration and amplitude) mapping tool in response to extrasystolic stimulation (S1-S2). Based on the results of our comprehensive test coverage, *KairoSight-3*.*0* can be employed to analyze complex cardiac optical datasets with accuracy and reliability.

## 2. METHODS

### 2.1. Animal Model

Animal protocols were approved by the Institutional Animal Care and Use Committee within the Children’s National Research Institute and followed the Nationals Institutes of Health’s *Guide for the Care and Use of Laboratory Animals*. Experiments were performed using adult, mixed sex Sprague Dawley rats (strain NTac: SD, from NIH Genetic Resource stock, Taconic Biosciences: Germantown New York) and adult, mixed sex Dunkin Hartley guinea pigs (GP). Animals were housed in conventional acrylic cages in the Research Animal Facility, under standard environmental conditions (12:12 hours light:dark cycle, 18-25º C, 30-70% humidity).

### 2.2. Isolated Heart Preparation and Langendorff-Perfusion

Animals were anesthetized with 2-3% isoflurane, until a surgical depth of anesthesia was reached as determined by a tail pinch test (rats) or ear pinch test (guinea pigs). Animals were euthanized by exsanguination following heart excision. The isolated, intact heart was submerged in ice cold cardioplegia and its aorta cannulated. The heart was then transferred to a temperature controlled (37ºC), constant pressure (70 mm/Hg) system for retrograde perfusion with modified Krebs-Henseleit buffer (mM: 118.0 NaCl, 3.3 KCl, 1.2 MgSO_4_, 1.2 KH_2_PO_4_, 24.0 NaHCO_3_, 10.0 glucose, 2.0 sodium pyruvate, 10.0 HEPES buffer, 2.0 CaCl_2_) continuously bubbled with carbogen (95% O_2_, 5% CO_2_). Intact heart preparations were allowed to equilibrate for 10-20 minutes on the perfusion system. Once the heart was stabilized, (-/-) blebbistatin (rat heart: 8.5 μM, guinea pig heart: 12.8 μM) was added to the perfusate to reduce metabolic demand and suppress contractile motion^17–19^.

### 2.3. Fluorescence Imaging

For optical CaT recordings, isolated hearts were loaded with a calcium indicator dye (Rhod2-AM: 100 μg rat heart, 150 μg guinea pig heart; AAT Bioquest, Sunnyvale CA). For optical AP recordings, isolated hearts were loaded with a potentiometric dye (RH237: 62 μg rat or guinea pig heart; AAT Bioquest). Fluorescent signals were acquired by illuminating the epicardial surface with a light emitting diode equipped with excitation filter (525 ± 25 nm; ThorLabs, Sterling VA). Emission filters were used to collect the calcium signal (585 ± 20 nm) or voltage signal (>700 nm)^4^. Image stacks were collected using high speed cameras (Dimax CS4: PCO-Tech, Kelheim Germany; Optical Mapping System from MappingLabs Ltd equipped with Prime BSI cameras, Teledyne Photometrix). Programmed electrical stimulation was implemented by placing an electrode (Harvard Biosciences, Holliston MA) on the epicardial surface (anteriorly positioned), which was controlled by an electrophysiology stimulator (Bloom Classic: Fisher Medical, Wheat Ridge CO; VCS-3001 stimulator: MappingLabs) set to 1 ms pulse width, 0.7-1.2 mA. Either a dynamic (S1-S1) or extrasystolic (S1-S2) pacing protocol was used, as indicated.

### 2.4. Optical Mapping and Analysis Software

Optical signals were analyzed using *KairoSight-3*.*0*, a customized semi-automated software developed in Python 3.8.5, which employs a graphical user interface (software and manual available: https://github.com/kairosight). In this study, we expanded the available features in *KairoSight-3*.*0* and validated the accuracy of its automated measurements with a user-directed algorithm developed in MATLAB. Briefly, an image stack (Tag Image File Format (.TIFF)) is uploaded to *KairoSight-3*.*0*, the image stack is then cropped and masked to generate a region of interest (ROI). Thereafter, signals within the ROI can be further processed by 1) spatial filtering, 2) temporal filtering, 3) drift removal, and 4) amplitude normalization. Next, the user selects a time window of interest (e.g., a single action potential) and the specified cardiac metric is mapped across the ROI (**Figure 1**). Newly implemented features in *KairoSight-3*.*0* include:

**Figure 1.**
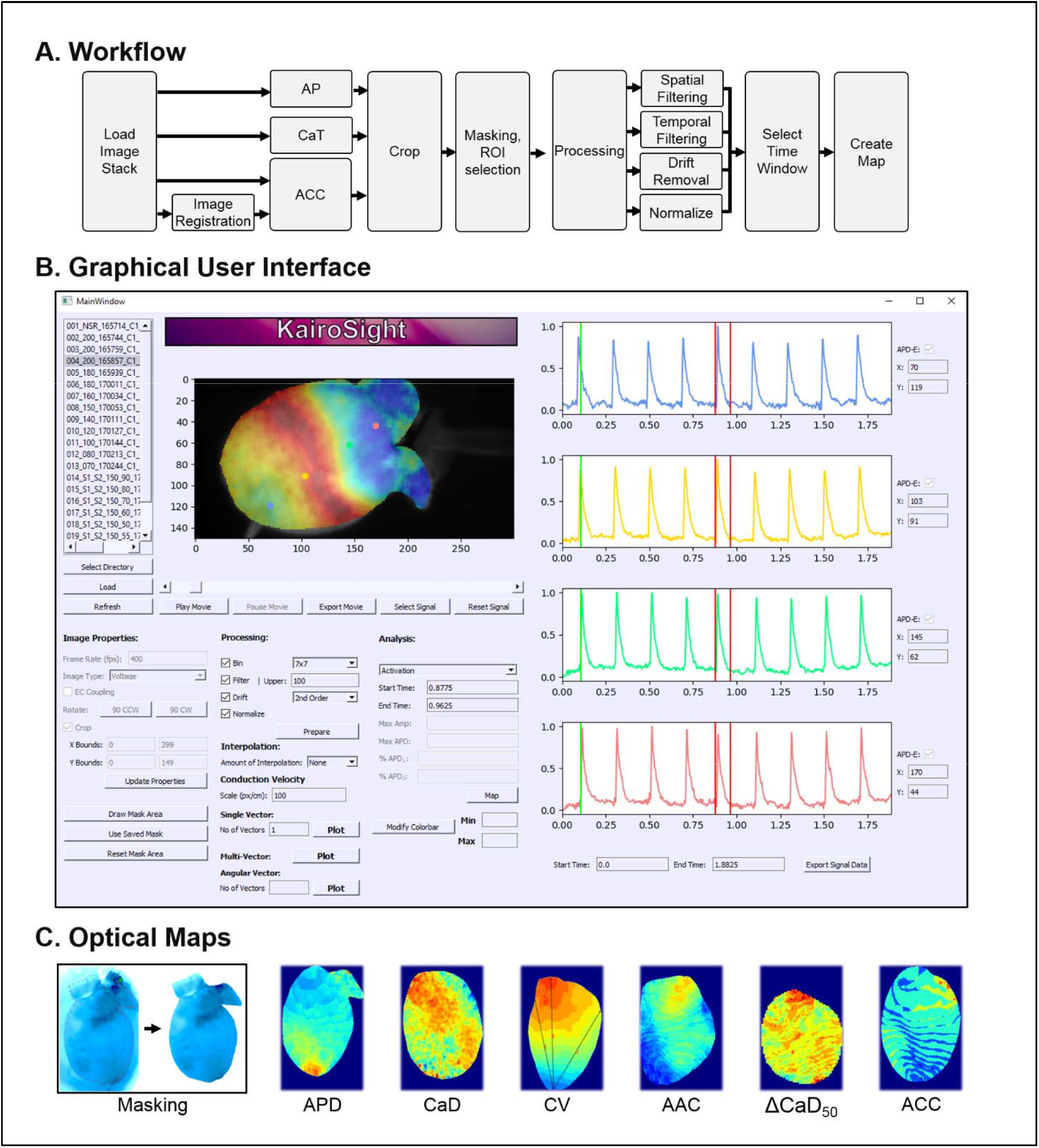
KairoSight-3.0 Overview. **A)** Illustration of *KairoSight-3.0* workflow. **B)** *KairoSight-3.0* graphical user interface. Action potentials on the right correspond to selected pixels (colored dots) on the epicardial surface. A time window of interest is selected (red bars) to highlight a single action potential. **C)** Example of *KairoSight-3.0* analysis features, including a masking tool to select a region of interest (left), and the generation of optical maps (right) corresponding to action potential duration (APD), calcium transient duration (CaD), conduction velocity (CV), action potential alternans coefficient (AAC), calcium coefficient from extrasystolic stimulation (ΔCaD_50_), action potential-calcium transient coupling (ACC).

#### Manual Masking Tool

This tool allows a user to manually select an ROI and assign a ‘zero’ value to negate any pixels that are located outside the defined ROI (**Figure 1C**). The user simply selects the desired ROI by mouse click, and the generated mask can then be applied to either a single image stack - or the mask can be saved and applied to additional image stacks from the same experiment. If the selected area is not satisfactory, the user can simply reset the mask.

#### AP Duration (APD) and CaT Duration (CaD)

*KarioSight* uses an updated algorithm to measure APD and CaD (**see Supplemental Figure 1**), which allows for a robust approximation of the offset point of the optical signal. Further, in the updated algorithm, optical signals are normalized within the selected time window (e.g., a single action potential) rather than across the entire image stack (e.g., multiple action potentials within the entire recording time).

#### Interpolation

This tool employs ‘Interp’ function in Python that performs linear interpolation for monotonically increasing sample points. The tool has been designed to avoid error due to low sample points within a particular time window of interest. For instance, when an APD is calculated at a certain percent repolarization, there may not be any sample point with in allowable error limit at the downstroke of an action potential. In such a case, using this linear interpolation tool with additional sample points allows the calculation to be obtained within the ‘null’ time window of interest (**Supplemental Figure 2**). Depending on the signal quality, a user can double or triple the interpolated sample points.

#### Signal-to-Noise Ratio (SNR)

This tool creates a pixel-to-pixel map of the SNR, calculated at each pixel as

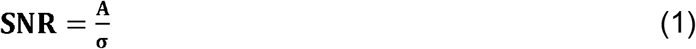

where “A” is the amplitude of the signal and “σ” is the standard deviation of noise, calculated at the baseline within the designated time window.

#### Conduction Velocity

This tool measures the scalar value of a velocity vector. The user can generate any number of conduction velocity vectors by defining the start and end point of each vector on a ‘pop up’ activation map (generated from the maximum upstroke rate).

#### Cardiac Alternans

This tool maps the degree of APD or CaT amplitude alternans, in cardiac preparations with consecutively alternating signals. The APD alternan coefficient (AAC) is calculated as

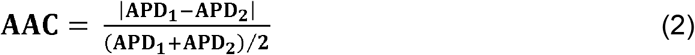

where “APD_1_” is the APD of the first action potential, “APD_2_” is the APD of the consecutive action potential. The APD can be examined at any specified repolarization value. The CaT alternan coefficient (CAC) is calculated as

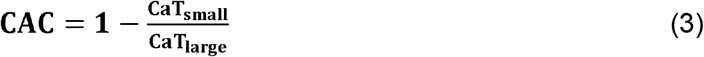

where “CaT_small_” is the amplitude of the smaller CaT, and “CaT_large_” is the amplitude of the larger CaT.

*KarioSight* includes three methods for mapping cardiac alternans: 1) fixed AAC, 2) dynamic AAC, and 3) user feedback AAC. Briefly, the fixed option allows a user to map AAC at any specified repolarization value. The dynamic option maps the AAC or CAC by automatically identifying the minima between the two consecutive peak signals within the alternating beats. The user feedback option allows the optimal level of repolarization to be determined from dynamic APD minima maps (**see Supplemental Figure 3-5**).

#### Extrasystolic Stimulation (S1-S2)

This tool generates maps corresponding to i) APD or CaD, ii) AP or CaT amplitude, iii) duration or amplitude coefficients when cardiac preparations are subjected to extrastimulus pacing (**see Supplemental Figure 6**). The APD coefficient (“Δ_APD_”) is calculated as

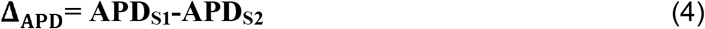

where “APD_S1_” is the APD from the S1 stimulus, “APD_S2_” is the APD from the S2 stimulus. The CaD coefficient (“Δ_CaD_”) is calculated as

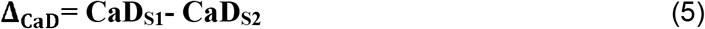

where “CaD_S1_” is the CaD from the S1 stimulus, “CaD_S2_” is the CaD from the S2 stimulus. The signal amplitude coefficient (“Δ_A_”) is defined as

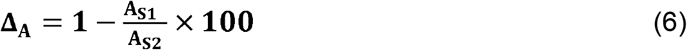

where A_S1_ is the amplitude of AP or CaT from stimulus S1, A_S2_ is the amplitude of AP or CaT from stimulus S2.

#### AP-CaT Coupling (ACC)

This tool can generate maps of AP-CaT coupling coefficient during activation (“ACC_act_”), defined by

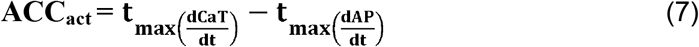

where 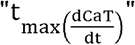 is the time at the maximum slope of the CaT upstroke and 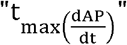 is the time at the maximum slope of the AP upstroke. This tool can also generate maps of AP-CaT coupling coefficient during repolarization (“ACC_rep_”), defined by

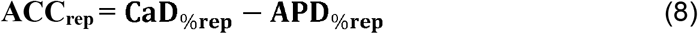

where “CaD_%rep_” is the CaD at a defined percent repolarization and “APD_%rep_” is the APD at a defined percent repolarization. Of importance, ACC mapping requires that voltage and calcium images are spatially aligned, which can be achieved by precise camera alignment and/or post-processing using image registration (**see Supplemental Figure 7**). In this study, image stacks were acquired using two aligned cameras and post-processing was also implemented using the overlay and translate tool in *Fiji software*^20^ (https://imagej.net/).

To validate the accuracy of automated measurements in *KairoSight-3*.*0*, a user-directed algorithm was developed in MATLAB (MathWorks) to construct manual maps (n=11). Three independent reviewers (FB, AR, and SA) marked the necessary fiducial points on each optical signal from every pixel (total of 1000-2000), and these derived values were then used to construct a manual map. The stack of voltage and calcium images underwent the same degree of post-processing for manual and automated analysis. The percent accuracy (“Ø_*x*_”) between manual and automated maps was defined by

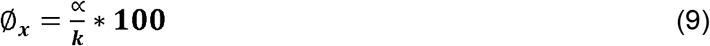

where “k” is the number of metrics (e.g., APD) in the ROI, and “∝“ is the number of metrics in the ROI that satisfy the following equation:

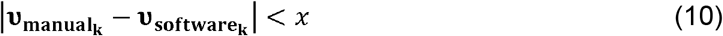

where 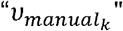 is the metric value calculated manually, 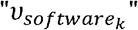 is the metric value calculated via software, and x is the maximum acceptable difference between the manual and software values (e.g., maximum duration difference of <5ms or <10ms, maximum alternan coefficient differences of <0.1, <0.15, or <0.2).

## 3. RESULTS

### 3.1. Validation of Automated *KairoSight-3*.*0* Software

To verify the accuracy of the newly implemented algorithms, optical maps were generated using two methods – user-specified manual mapping or *KairoSight-3*.*0* automated mapping – and a total of 11 cardiac metrics were analyzed (**Figure 2**). First, we analyzed duration measurements by creating manual maps of APD_30_, APD_80_, CaD_30_, and CaD_80_ (**Figure 2A**). Next, we analyzed the same image stacks using *KairoSight-3*.*0* to create software maps (**Figure 2B**). Finally, manual and software maps were quantitatively compared (**Figure 2C**). We observed excellent correlation between manual and software-generated maps; for example, both approaches resulted in APD_80_ measurements that varied between 30-55 ms with longer APD values located near the left ventricular apex. Manual and software maps for APD and CaD suggest high accuracy, with >97% of manual and software values falling within 10 ms of each other and >75% within 5 ms (**Figure 2C**: ∅_*x*_ ≥98% when x<10 ms; ∅_*x*_ ≥76% when x<5 ms, n=8 maps, 1000-2000 pixels). We also analyzed cardiac alternans using three different tools: APD alternans at a specified repolarization value (e.g., APD_50_), APD alternans measured dynamically at the minima between two peaks, and calcium amplitude alternans (**Figure 2D-I**). Each of these cardiac metrics yielded similar spatial maps, whether generated manually or via *KairoSight-3*.*0* automated mapping. As an example, dynamic alternan coefficient maps demonstrate high accuracy with >93% of AAC and CaC coefficient values differing by less than 0.2 (**Figure 2I**: ∅_*x*_ ≥94% when x<0.2; ∅_*x*_ ≥78% when x<0.2; ∅_*x*_ ≥74% when x<0.1, n=3 maps, 1000-2000 pixels). Given the large number of pixels analyzed (1000-2000), some degree of variability is expected due to user error in marking fiducial points and/or noisy optical signals near the edges of the tissue preparation. To the best of our knowledge, our study is the first to validate cardiac analysis algorithms with a user-specified manual mapping approach.

**Figure 2.**
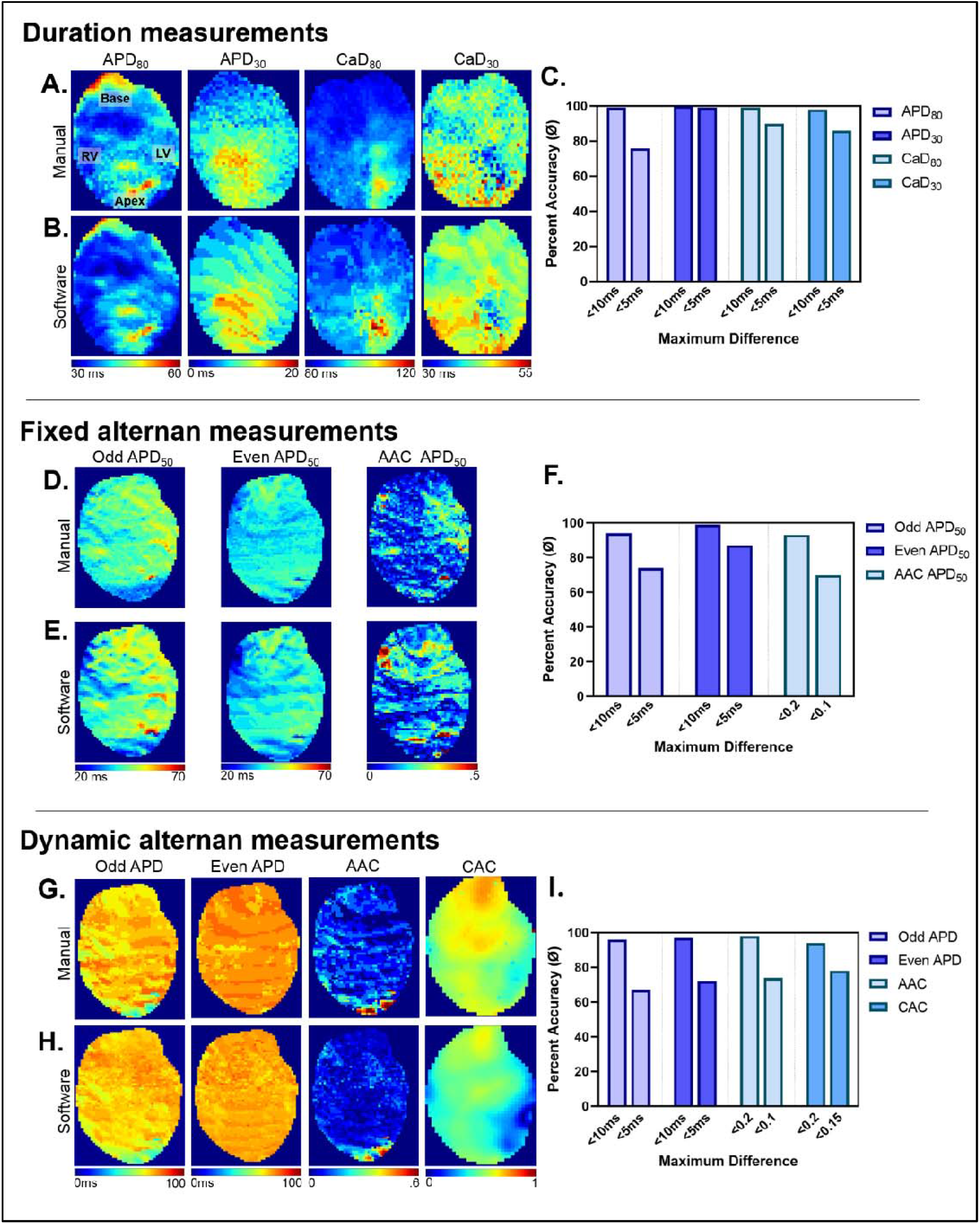
Validation of Semi-Automated Software Analysis. Optical signals were analyzed using *KairoSight*-3.0, and the accuracy was compared with manual measurements. **A-C)** Maps of APD and CaD derived manually, using *KairoSight-3.0* software, and the percent accuracy of the automated measurement. **D-F)** Analysis of alternans from sequential APD measurements (odd, even beats) at a specified repolarization phase (50%, “fixed”), with corresponding coefficient measurement (AAC). Results derived manually, using *KairoSight-3.0* software, and the percent accuracy of the automated measurement. **G-I)** Analysis of alternans from sequential APD measurements (odd, even beats) at the maximum detected repolarization phase between consecutive beats (“dynamic”), using manually analysis, *KairoSight-3.0* software, and the percent accuracy of the automated measurement. Note the longer duration time compared to “fixed” approach indicates repolarization >50%. An example of calcium alternan coefficient (CAC) map is also shown. Optical maps derived from an intact, rat heart loaded with a voltage- or calcium-sensitive dye.

### 3.2. Impact of Signal Quality on Data Analysis

Cardiac optical mapping experiments are highly dependent on fluorescence signal quality, which can vary between replicate experiments based on dye loading and washout, photobleaching, illumination intensity and exposure time^21–23^. As such, spatial and temporal filtering are frequently applied to improve signal quality^9,10,24^. Moreover, specific tissue regions can have more noise compared to other regions, particularly areas with shadows, suboptimal light illumination, or edges of the cardiac tissue preparation^25^. Accordingly, SNR maps are a useful tool to quickly identify tissue regions with more or less noise, with the latter likely to yield more reliable optical signals.

To demonstrate the use of the SNR mapping tool in *KairoSight-3*.*0*, we modulated the excitation light intensity for a set of imaging experiments, and then applied spatial and/or temporal filtering to the acquired optical data sets (**Figure 3**). In the raw optical signals, voltage SNR decreased from ∼30 at 100% light intensity to ∼5 at 40% light intensity (**Figure 3A**). As expected, SNR was significantly improved by processing the image stack, through spatial and temporal filtering in *KairoSight-3*.*0*. Despite improvements in SNR, the signal quality can still vary across the epicardial surface, as indicated by a heterogeneous SNR map. Further, we also tested the impact of signal quality on cardiac metric measurements. In the raw optical signals, CV was easily measured in datasets with high SNR at maximal light intensity (e.g., LED 100%), but CV measurements became unreliable in datasets with low light intensity (e.g, LED 40-70%; **Figure 3B**). From the processed signals, CV was more easily measured in datasets with maximal-to-moderate light intensity (e.g., LED 100-60%) – even within tissue regions that had a relatively low SNR (∼8-10 at 60% light intensity). Similarly, in both the raw and processed maps, APD_30_ and APD_80_ measurements were comparable between the raw and filtered data at the maximal light intensity (e.g., LED 100%) (**Figure 3C, D**). But, with lower light intensity (and reduced SNR), the raw APD maps were less interpretable with highly heterogeneous APD distribution observed within a relatively small pixel area. Collectively, SNR mapping in *KairoSight-3*.*0* – either with or without processing – can be a highly useful tool for identifying regions of interest with superior signal quality and/or identifying poorly labeled tissue regions that might need to be excluded from downstream analysis.

**Figure 3.**
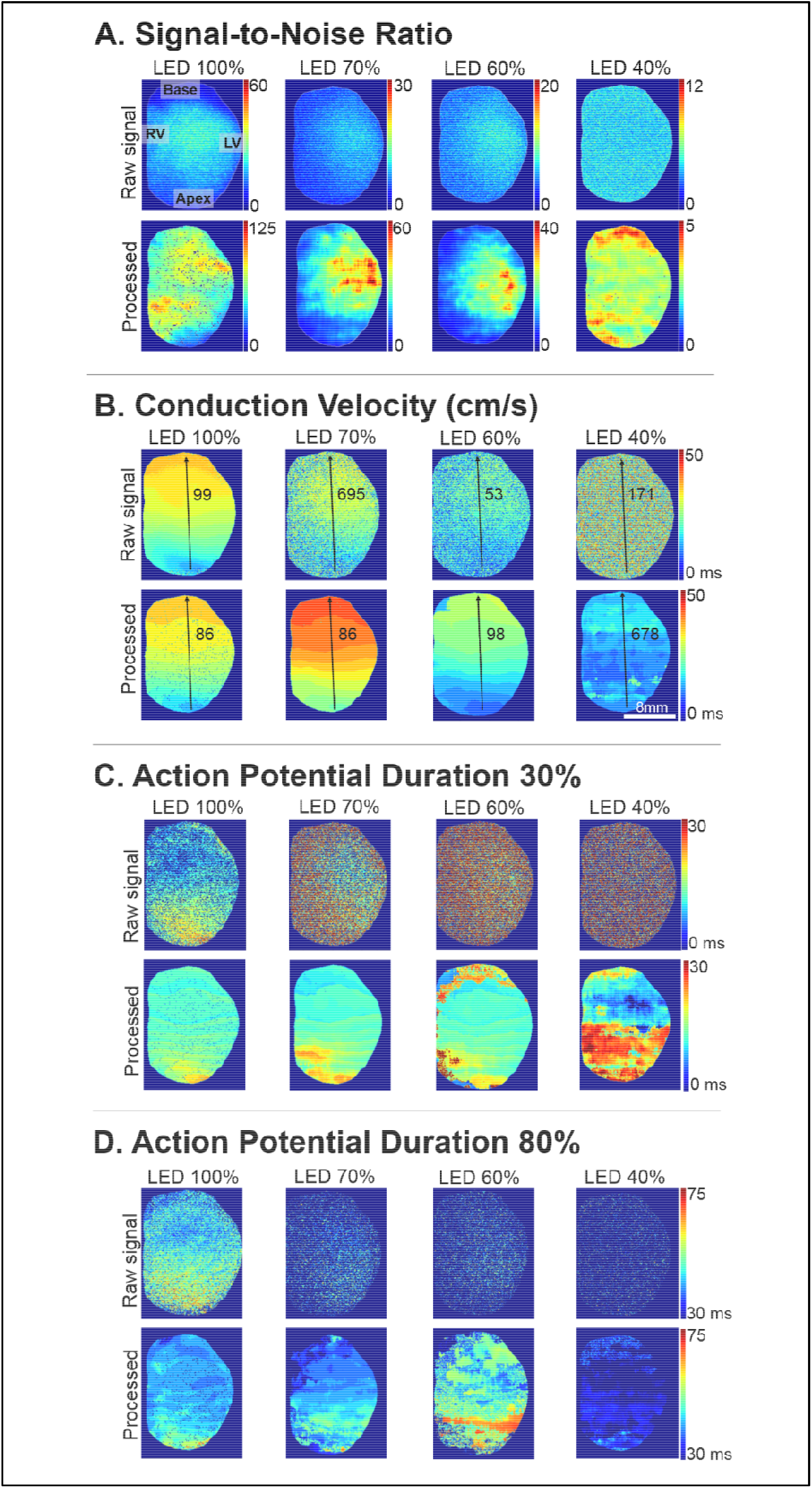
Impact of Signal Quality and Filtering on Data Analysis. **A)** Reducing the excitation light intensity (e.g., 40-100%) results in a lower signal-to-noise ratio (note difference in scale). Post-processing can be applied (e.g., temporal filter, spatial averaging) to aid in the measurement of **B)** conduction velocity, **C)** APD at 30% repolarization, and **D)** APD at 80% repolarization. Example of optical maps derived from an intact, rat heart loaded with a voltage-sensitive dye.

### 3.3. Analyzing Cardiac Alternans in *KairoSight-3*.*0*

Cardiac alternans are broadly defined as beat-to-beat oscillations in electrical activity (APD alternans), intracellular calcium cycling (calcium alternans), or mechanical activity, and the magnitude of these sequential oscillations have been linked to cardiac arrhythmias^26,27^. Accordingly, the action potential or calcium transient alternan coefficient maps are a useful tool for visually identifying and classifying concordant and/or discordant alternans, which are more commonly observed with faster pacing, low temperature, cardiac disease or injury^28–33^. To test the utility of the AAC and CAC mapping feature in *KairoSight-3*.*0*, optical data sets were acquired from the anterior epicardial surface of a Langendorff-perfused guinea pig heart in response to slower (200 ms PCL) and faster pacing rates (100 ms PCL; **Figure 4**). At the slower pacing rate, maps generated using the fixed AAC tool between 20-60% repolarization yield minimal differences – indicating the lack of APD alternans (**Figure 4A**). Similar results are observed for CaT amplitude measurements at the slower pacing rate, with a CAC value <0.15 between two consecutive beats (**Figure 4C**). However, at the faster pacing rate, fixed AAC maps indicate alternans at all measured intervals between 20-60% repolarization (**Figure 4A**). In this example, alternans were present at the left ventricular base with maximal difference between sequential beats (AAC > 0.5), but the lateral-apical region had minimal difference between beats (AAC < 0.1). Similar regional heterogeneity was observed with calcium alternans at the faster pacing rate, with a small CAC at the apex (<0.15) and greater CAC at the left ventricular base (>0.5). Dynamic AAC maps (**Figure 4B**), which uses the minima between two signal peaks of alternating beats showed even more pronounced dispersion between the apical region (<0.1) and the ventricular base (>0.65).

**Figure 4.**
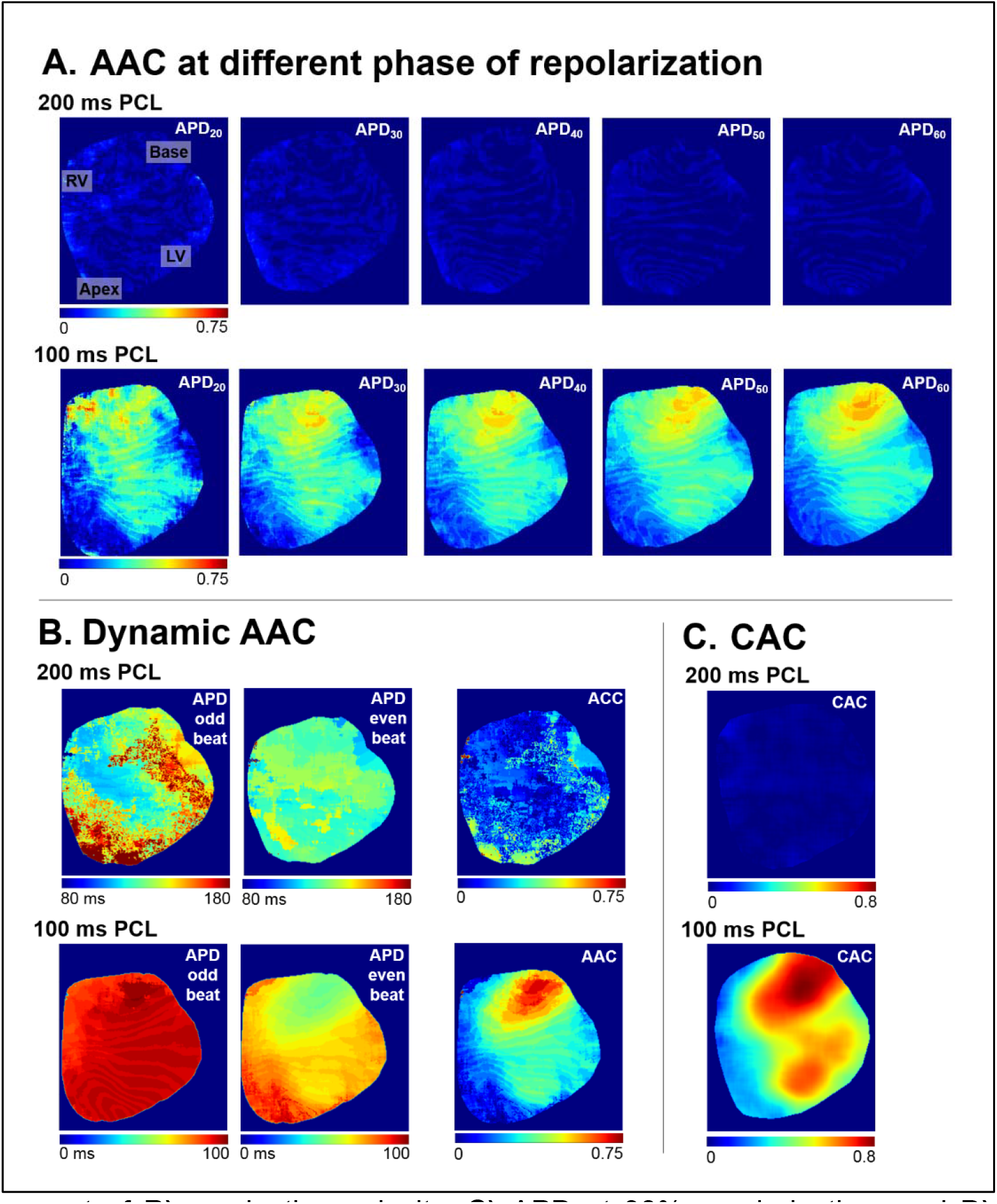
Mapping of Cardiac Alternans. **A)** AAC maps generated from two sequential action potentials using the “fixed” alternan tool at varying phases of repolarization (20-60% APD). Top: AAC at a slower pacing rate (200 ms PCL); Bottom: AAC at faster pacing rate (100 ms PCL). **B)** APD measurements for sequential (odd, even) action potentials and the calculated AAC using the “dynamic” alternan tool. Variables measured at a slower (200 ms PCL) and faster rate (100 ms PCL). **C)** CAC maps generated from two sequential calcium transients using the “dynamic alternans” tool. CAC derived at a slower (200 ms PCL) and faster rate (100 ms PCL). Optical maps derived from an intact, guinea pig heart loaded with a voltage- or calcium-sensitive dye. AAC = AP alternans coefficient, CAC = calcium alternans coefficient.

### 3.4 Analyzing Cardiac Electrophysiology Metrics in Response to Extrasystolic Stimulation

Extrasystolic (S1-S2) stimulation is frequently employed in cardiac electrophysiology studies to characterize tissue refractoriness, assess electrical restitution, interrogate arrhythmia susceptibility, and/or avoid myocardial ischemia that can result from fast dynamic pacing (S1-S1)^1,3,34,35^. Accordingly, we implemented an S1-S2 mapping tool in *KairoSight-3*.*0* to quantify cardiac metrics during the S1 beat, S2 beat, and measure the corresponding difference. In a representative example (**Figure 5**), an intact guinea pig heart was paced from the left ventricular apex and the S1-S2 interval was decremented stepwise by 10 ms (200 ms – 110 ms). Maximum APD_50_ shortened from 100 ms at 200 ms PCL to 70 ms at 140 ms PCL, with a heterogeneous Δ_APD50_ that varied between 10-20 ms without discernable regional specificity (**Figure 5A**). Similarly, CaD_50_ shortened from 105 ms to 75 ms (200 vs 140 ms PCL, respectively), with a slightly greater Δ_CaD50_ observed near the left ventricular base (**Figure 5B**). Further, a striking apico-basal heterogeneity pattern was observed in the S2 CaT amplitude – with larger amplitudes observed at the apex/mid region, and shorter amplitudes at the basal region (**Figure 5C**).

**Figure 5.**
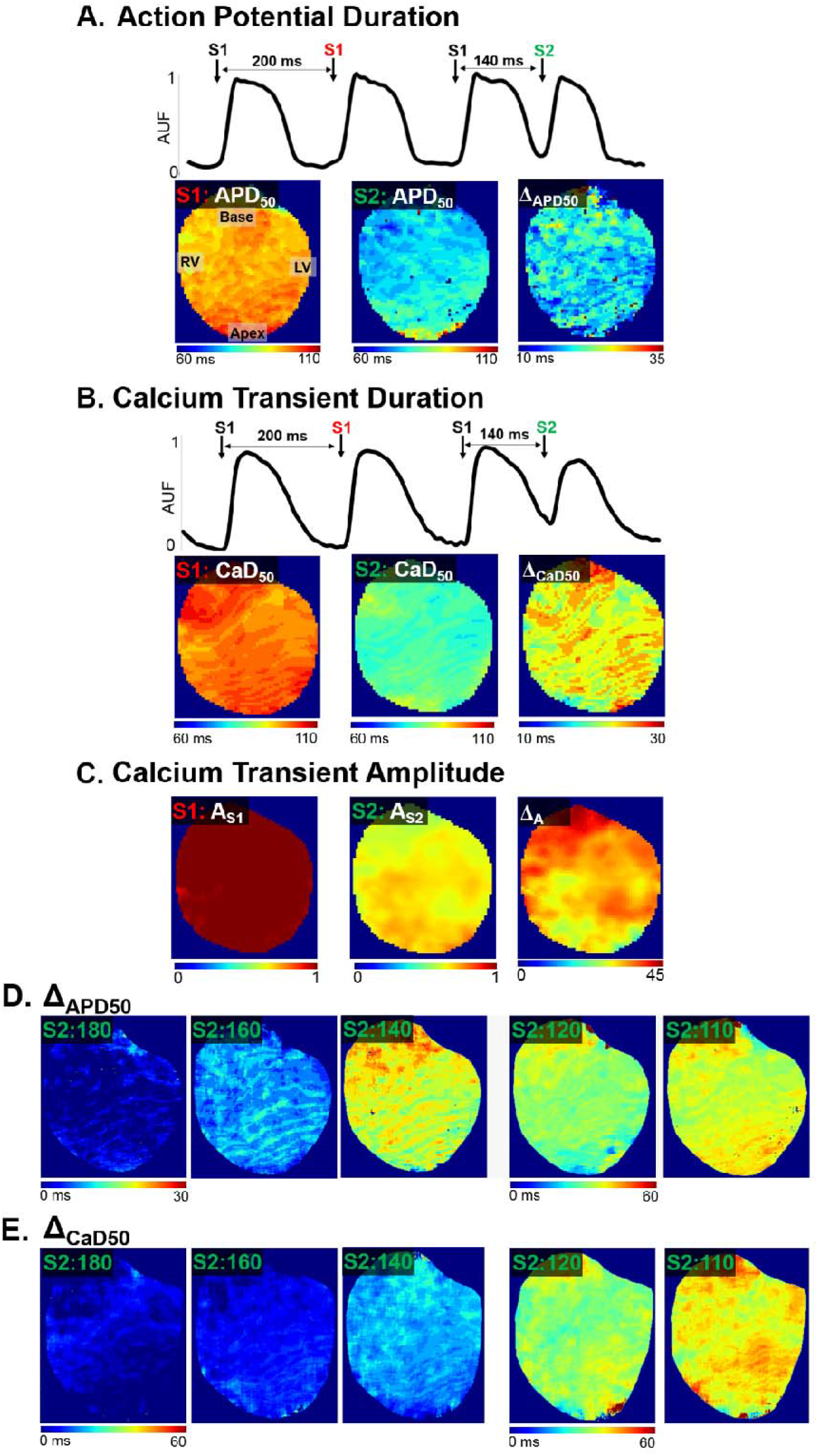
Effect of Extrasystolic Stimulation on Cardiac Metrics. **A)** Top: Example of optical action potential recordings resulting from regular (S1: 200 ms PCL) and extrasystolic stimulation (S2: 140 ms PCL). Bottom: Corresponding optical maps for APD_50_ during S1, S2, and the map of the resulting difference. **B)** Top: Example of optical action potential recordings resulting from regular (S1: 200 ms PCL) and extrasystolic stimulation (S2: 140 ms PCL). Bottom: Corresponding optical maps for CaD_50_ during S1, S2, and the map of the resulting difference. **C)** Corresponding optical maps for CaT amplitude during S1, S2, and the resulting difference between the two maps. **D)** Difference maps of APD_50_ as S2 PCL is decremented (180-110 ms). Note different scale bar at faster frequencies. **E)** Difference maps of CaD_50_ as S2 PCL is decremented (180-110 ms). Fluorescent signals collected from guinea pig hearts.

### 3.5 Epicardial conduction velocity measurements

Epicardial conduction velocity measurements are commonly measured to track changes in inward sodium current, cell-cell electrical coupling, interstitial volume, and/or structural remodeling that can hinder action potential propagation, and may create a substrate for arrhythmia development^36–39^. During sinus rhythm, the wavefront of the propagating electrical impulses create random breakthrough patterns in the epicardial surface; as such, epicardial conduction velocity measurements are acquired with external pacing. The measurements of conduction velocity can be affected by the location of the pacing site^40,41^, given the presence of intraventricular regional variations in fiber orientation and ion channel expression^42–44^. Accordingly, we implemented a CV measuring tool in *KairoSight-3*.*0* and provide examples of semi-automated CV measurements based on pacing site (e.g., mid-epicardium, apex, left or right ventricular base, **Figure 6**). Directional heterogeneity in the impulse propagation is most apparent when the heart is paced near the septum or mid ventricular region (**Figure 6A**, 58 cm/s longitudinal CV, 33 cm/s transverse CV). Apical/basal pacing provides robust apico-basal CV measurements, but it is difficult to discern transverse:longitudinal CV values as the elliptical wavefront is partially concealed from view (**Figure 6B-D**). Notably, *KairoSight-3*.*0* allows the user to select multiple vectors for a comprehensive analysis of epicardial CV.

**Figure 6.**
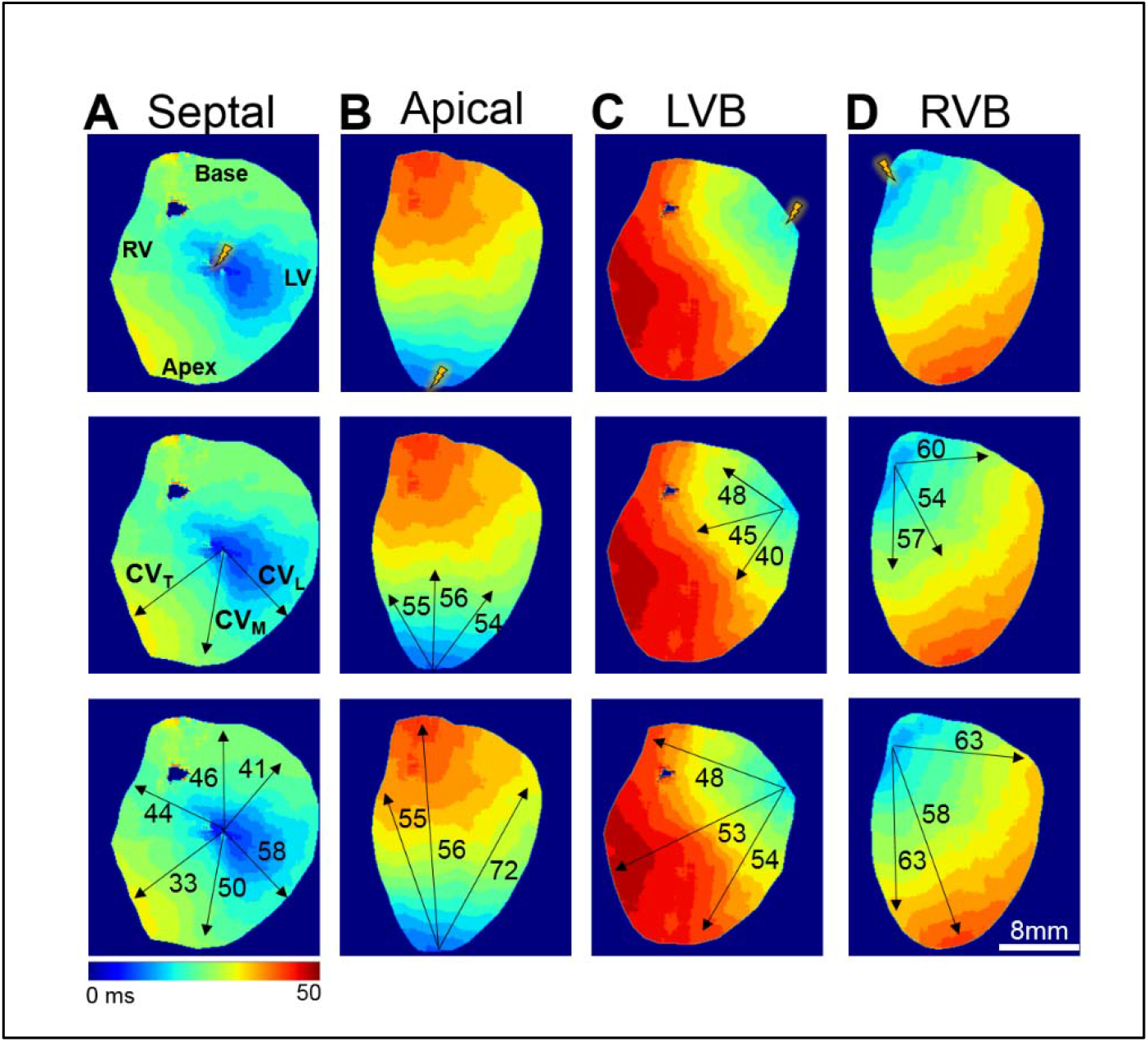
Conduction Velocity Measurements According to Pacing Location. **A)** Top: Illustration of septal pacing site. Middle: Examples of electrical propagation in the mid, longitudinal, and transverse directions. Bottom: Example conduction velocity measurements collected from six vectors. **B)** Top: Illustration of apical pacing site. Middle: Example conduction velocity measurements half-way across the epicardial surface. Bottom: Example conduction velocity measurements from apex to base. **C)** Top: Illustration of pacing site at the left ventricular base (LVB). Middle: Example conduction velocity measurements half-way across the epicardial surface. Bottom: Example conduction velocity measurements across the entire heart. **D)** Top: Illustration of pacing site at the right ventricular base (RVB). Middle: Example conduction velocity measurements half-way across the epicardial surface. Bottom: Example conduction velocity measurements across the entire heart. Scale bar denotes activation time; fluorescent signals collected from rat hearts.

### 3.6 Action potential-calcium transient coupling

Simultaneous mapping of voltage and calcium signals can yield important information on cardiac excitation-contraction coupling, the process by which an electrical excitation is linked to contraction. Alterations in excitation-contraction coupling can impact APD, initiate after-depolarizations, and trigger electrical/mechanical alternans that can be proarrhythmic^45,46^. As such, *KairoSight-3*.*0* has the necessary tools to generate maps of AP-CaT coupling (ACC) coefficient during activation and repolarization from co-located optical signals. In **Figure 7**, we demonstrate ACC coefficient mapping using spatially-aligned images collected from a guinea pig heart at two different pacing rates (200 ms, 120 ms PCL). At the slower pacing rate, the ACC activation (ACC_act_) map indicates a very short delay between AP and CaT activation (∼1 ms) at the apical region and a slightly longer delay near the base (∼2-3 ms, **Figure 7B**). At the faster pacing rate, the delay between the AP and CaT upstroke is more evident and regional variability persists with a longer ACC_act_ at the base of the ventricle. We also generated ACC repolarization (ACC_rep_) maps at different phases of recovery (**Figure 7C**); at the slower pacing rate ACC_rep_ is mostly negative at the mid and basal regions during early repolarization (30-50%) indicating that the CaD is shorter, but ACC_rep_ becomes progressively more positive during later phases of repolarization (60-80%). However, this apico-basal heterogeneity is less evident at faster pacing rates (120 ms PCL). In this example, the only regional heterogeneity is in the early phases of repolarization (30-40%) between the right lateral (−5 to -8 ms) and left lateral (5-10 ms) regions of the ventricle.

**Figure 7.**
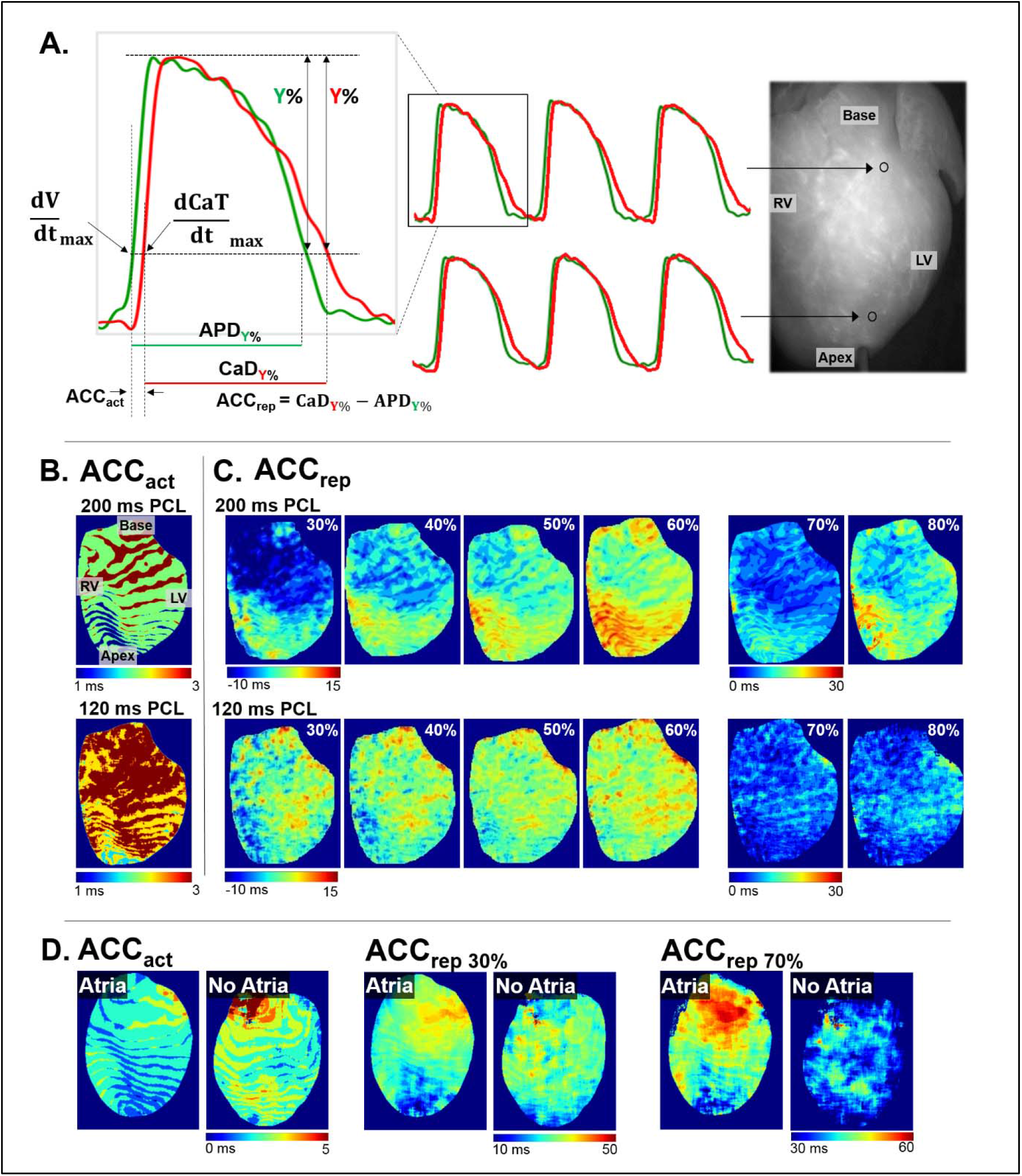
Action Potential-Calcium Transient Coupling (ACC). **A)** Illustration of ACC measurements, with overlay of optical action potentials (green) and intracellular calcium transients (red). **B)** ACC_act_ maps generated in response to slow (200 ms PCL) or fast pacing rate (120 ms PCL). **C)** Multiple ACC_rep_ maps derived at different phases of repolarization (30-80%), in response to slow (200 ms PCL) or fast pacing rate (120 ms PCL). ACC_act_ = action potential-calcium transient coupling during activation, ACC_rep_ = action potential-calcium transient coupling during repolarization. **D)** Effect of atrial removal on action potential-calcium transient coupling during activation (ACC_act_) and repolarization (ACC_rep30%_, ACC_rep70%_)

Optical mapping studies often use external stimulation to quantify AP and CaT metrics, given the rate-dependency. However, to prevent competitive stimulation via the sinus node, the atria may be removed from heart preparations to allow for external pacing at slower rates. As an example, we demonstrate the effect of atrial removal on cardiac AP-CaT coupling (**Figure 7D**). ACC_act_ map had an apico-basal gradient in the intact heart, which shifted to a more lateral gradient after atrial removal with a longer AP-CaT delay in the right ventricle. A similar trend was observed for ACC_rep_ maps at 30-70% with a clear apico-basal gradient when the atria were intact, but this gradient was absent after the atria were removed. For Langendorff studies, atrial dissection should be performed with careful consideration, as removal can perturb normal physiology (e.g., atrial natriuretic peptide release) and has previously been shown to alter myocardial response to pharmacological agents^47–49^.

## 4. DISCUSSION

In the presented article, we report additional and updated features of our previously developed cardiac optical mapping software *KairoSight*^16^. To verify the accuracy of the newly implemented algorithms, optical maps were generated from user-specified manual mapping and *KairoSight-3*.*0* automated mapping, and the accuracy between the two methods were compared with 11 different cardiac metrics. This study may be the first to validate optical cardiac maps using a manual mapping approach (1000-2000 pixels analyzed). Our validation study suggested reasonable accuracy in the software measurements (67% ≤ ∅_*x*_ ≤ 96%). Moreover, this updated software includes many additional features that can support research queries related to signal quality assessment, cardiac electrophysiology and intracellular calcium handling, including: 1) SNR mapping, 2) analysis of cardiac alternans (fixed, dynamic, user-specified tools), 3) conduction velocity measurements applying using user-specified vectors, 4) ACC mapping, and 5) metric mapping in response to extrasystolic stimulation. Further, we highlight the impact of variable experimental techniques (e.g., pacing site location, atrial removal) on cardiac optical mapping.

### 4.1. Quantitatively validated cardiac optical mapping software

In recent years, the cardiac electrophysiology research field has benefitted from a number of software options to support the analysis and mapping of cardiac imaging data^11–14,16^. However, the accuracy of these software tools has been largely limited to comparisons between normal and arrhythmogenic hearts, and to the best of our knowledge, this study is the first to quantitatively validate automated software measurements with a user-specified manual approach. The robustness of automated measurements is important, particularly for duration measurements, when there can be uncertainty in identifying a specific fiducial point at the signal downstroke due to experimental variables (e.g., slower frame rate, low SNR, slight motion artifacts from contracting tissue). Our revised algorithm optimizes the offset in fiducial point identification by allowing an error window and adding linearly interpolated values, as required (**see Supplemental Figure 2**). The result of this quantitative validation supports the robustness of the revised algorithm in automated duration calculations. Furthermore, manual mapping was performed by 3 independent reviewers (FB, AR, SA), which minimized reviewer bias. Slight discrepancies between automated versus manual mapping are expected due to human error in identifying fiducial markers, and or low SNR particularly at the edges of the cardiac preparation.

### 4.2. Novel cardiac metric mapping modules

#### 4.2.1. SNR mapping

Optical mapping experiments are highly dependent on fluorescent signal quality, which can vary between experiments based on uniform dye loading, dye internalization, dye washout, photobleaching, insufficient illumination or the presence of shadows. Accordingly, areas of low SNR can interfere with cardiac metric measurements. In this updated software, we include an SNR mapping feature to quickly identify signal quality and inform downstream signal analysis. Our software application also includes spatial and frequency filters that can be applied to improve signal quality.

#### 4.2.2. Mapping during dynamic and extrasystolic stimulation

One of the newly added features in *KairoSight-3.0* allows for accurate cardiac metric mapping, even at fast rates when fusion can occur between two consecutive signals. For example, fused optical signals can be observed during AP or calcium alternans, which has been mechanistically linked to arrhythmogenesis. Similarly, fused beats can be observed during extrasystolic stimulation (S1-S2), which is frequently employed to determine tissue refractoriness, characterize restitution, or as an alternative to dynamic pacing to minimize tissue ischemia. Analyzing fused signals is computationally challenging, with difficulty arising in identifying key fiducial points. Our validated APD and CaT alternan maps using either fixed or dynamic mapping tools shows robustness in the algorithms. While the ‘fixed alternan’ tool allows the use to specify the signal downstroke magnitude (e.g., % repolarization), the ‘dynamic alternan’ tool allows for automated identification of the fiducial point (midpoint) at which the maximum alternans may occur (**see Supplemental Figure 4**). This novel feature allows robust mapping of discordant alternans, which are characterized by spatial heterogeneity. Further, the software can also map S2 APD, AP amplitude, CaD, or CaT amplitude and the corresponding coefficients, which again demonstrates usability in the analysis of fused beats.

#### 4.2.3. AP-CaT coupling

Studies report AP-CaT activation delay or repolarization latency (often reported as a phase plot, AP-CaT delay trace) to provide information on excitation-contraction coupling ^5,50,51^. *KairoSight-3.0* allows for robust AP-CaT coupling mapping across the surface of the cardiac preparation – during both activation and repolarization. This tool can be used to characterize spatial heterogeneity (base vs apex, left vs right ventricle) in ACC, which is influenced by rate dependency of key transmembrane ionic currents (*I*_Na_, *I*_to_, *I*_Kr_, *I*_Ks_), calcium release channels (RyR) and transporters (SERCA pump, Na-Ca^2+^ exchanger).

#### 4.2.4. Experimental variables

Optical mapping experiments are most frequently performed on excised, intact heart preparations – although laboratories may use different techniques depending on the experimental question being investigated. We highlight two such examples (e.g., pacing site location, atrial removal) to demonstrate how different approaches can impact optical mapping. We utilized 3-6 vectors in *KairoSight-3.0* to measure epicardial CV in response to external pacing and varied the pacing location. Our data illustrates that CV measurements can vary, even within the same heart, based on pacing near the septum, base, or apex – and that the degree of variation depends on the vector direction in respect to the activation wavefront. The dependency of CV on pacing electrode location can be explained by the fact that expression of Na+ channels, connexin proteins and non-conducting tissue is heterogeneous across the epicardium, which impacts the activation propagation. We also performed a set of imaging experiments, in which cardiac metrics were quantified before and after atrial removal (the latter is often performed to avoid competitive activation from the sinus node, which can interfere with external pacing). Our results suggest that atrial removal impacts ACC_act_ and ACC_rep_ – with distinct heterogeneity gradients before and after atrial removal. Therefore, care should be taken to interpret optical maps with consideration of the technical variables used in each experiment.

## 5. CONCLUSION

We present an updated version of our previously developed software package, *KairoSight-3.0* with enhanced features for cardiac optical mapping and analysis. Through independent manual validation of optical maps, our results suggest robust accuracy of the automated measurements. The newly incorporated SNR mapping feature can streamline data analysis by identifying tissue regions with poor signal quality. Other newly incorporated features (e.g., action potential-calcium transient coupling, alternans mapping, conduction velocity measurements) allow for novel electrophysiology metric mapping, which can prove useful for characterizing spatially heterogeneous events in normal and arrhythmogenic hearts.

## 6. ACKNOWLEDGEMENTS

The authors gratefully acknowledge Blake Cooper for assistance with optical mapping experiments.

## 7. FUNDING SOURCES

This work was supported by the National Heart, Lung, and Blood Institute (R01HL139472, R01HD108839), Children’s National Research Institute, Children’s National Heart Institute, and the Gloria and Steven Seelig family.

## Abbreviations

AAC: APD alternans coefficient
ACC: Action potential – calcium transient coupling coefficient
AP: Action potential
APD: Action potential duration
CAC: CaT alternans coefficient
CaT: Calcium transient
CaD: Calcium transient duration
CV: Conduction velocity
EP: Electrophysiology
ROI: Region of interest
SNR: Signal-to-noise ratio

## Notes

### Competing Interest Statement

The authors have declared no competing interest.

