## Supplemental File for "*KairoSight-3.0*: A Validated Optical Mapping Software to Characterize Cardiac Electrophysiology, Excitation-Contraction Coupling, and Alternans"

**Software for the Analysis Cardiac Electrophysiology and Excitation-Contraction Coupling Data with Enhanced Features and User-Specified Validation**

Kazi T Haq^1,2^, *Anysja Roberts^1,2^, *Fiona Berk^1,3^, Samuel Allen^1^, Luther Swift^1,2^, Nikki Gillum Posnack^1,2,4^

**1. APD and CaD mapping:** Figure 1 illustrates how the *KairoSight-3.*0 algorithm approximates AP or CaT duration. A duration (APD or CaD) is given by,

$$\Delta t=t_{on}-t_{off}$$

where, t_on_ = time point at ${\frac{dV}{dt}}_{max}$ or ${\frac{d{Ca}_{i}}{dt}}_{max}$

t_off_ = time point at desired % of repolarization/Ca_i_ reuptake.

To find t_off_ (inset) the algorithm searches for all the time point at a certain x% (30% in the Figure) within a searching window Y (x ± 5). This is done because at exact x% there may not be any sample point. In the next step, the algorithm identifies t_off_ as the time point corresponding to a point that is nearest to x. In this example in the figure, two points (o, o’) fall within Y and the algorithm would find t_off_ at time point corresponds to o’, since b<a.

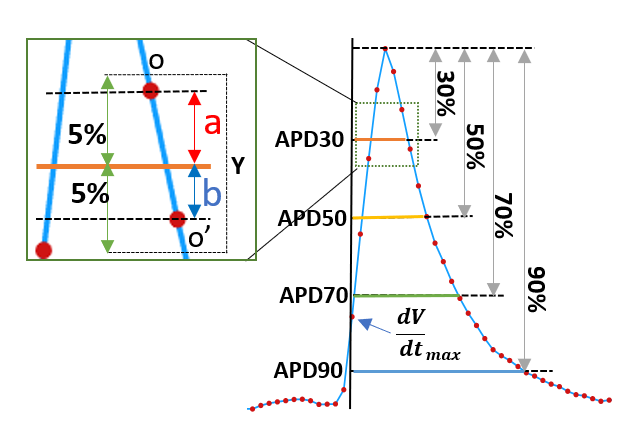

**Figure 1.** Example of APD calculations.

**2. Interpolation:** As shown in Figure 2 (top panel) linear interpolation increases intermediate sample points in the AP trace (green dots), compared with the original (red dots). The bottom panel shows the effect of interpolation on APD70 mapping. The map with the original sample points resulted in negative and ‘NaN’ numbers in the calculation as shown in the bottom panel. In this example, the duration calculating algorithm did not find a point at 70±5%, which is reflected by the negative or ‘NaN’ values in the map (left).

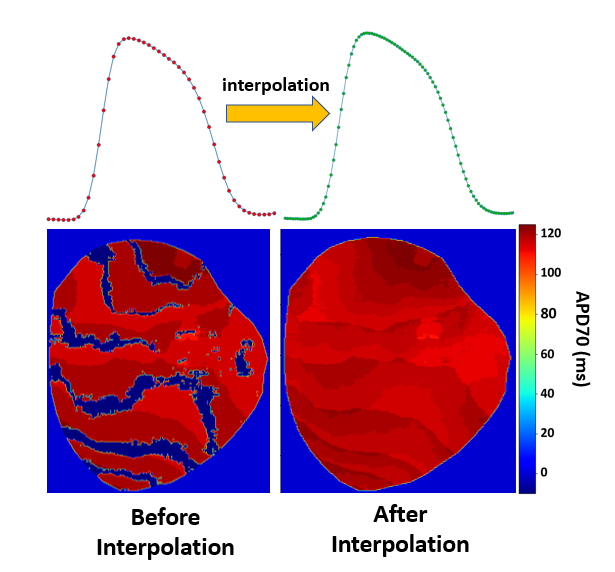

**Figure 2.** Example of interpolation

**3. Alternans**

A. Fixed AAC

As shown in Figure 3, user can choose any repolarization phase to map AAC. Note that the ‘fixed’ option is not applicable for CAC mapping, which is generally a CaT amplitude-based measurement.

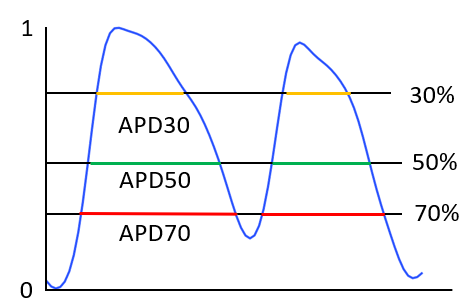

**Figure 3.** Example of “fixed” option for AP alternan measurements.

B. Dynamic AAC and CAC

Unlike the fixed AAC tool, dynamic AAC and CAC measurements do not require user input. It is ‘dynamic’ because it measures the parameters of the metric dynamically as much as a signal will allow (depending on degree of recovery). In the event when the alternans are discordant in nature, there could be two possible scenarios (Figure 4A and 4B). As shown in Figure 4A, the ‘mid-point’ (M) is the maximum repolarization phase of beat1 before beat2 depolarization occurs. However, beat2 repolarization approximately ends at point E. In this scenario,

Y1 > Y2

where, Y1 = amplitude of M

Y2 = amplitude of E

The dotted line D, drawn at M, intersects the downstroke phase of beat2 is the maximum repolarization (reuptake for a CaT signal) phase that can be calculated. The end point of APD2 is calculated in the intersection (X). In scenario 2, Y2>Y1 (Figure 4B). In this case, line D drawn from E, and which intersects downstroke line of beat2 at X, and end point of APD2 is calculated at X. To calculate dynamic AAC and CAC, first the algorithm identifies M and E, and next it calculates Y1 and Y2. Whichever of the Y1/Y2 relationships a signal satisfies, the algorithm calculates the APD/CaT amplitudes accordingly. Note that the algorithm is used to identify midpoints (M) between the two consecutive beats for a duration calculation, is also employed to calculate amplitude-based CAC.

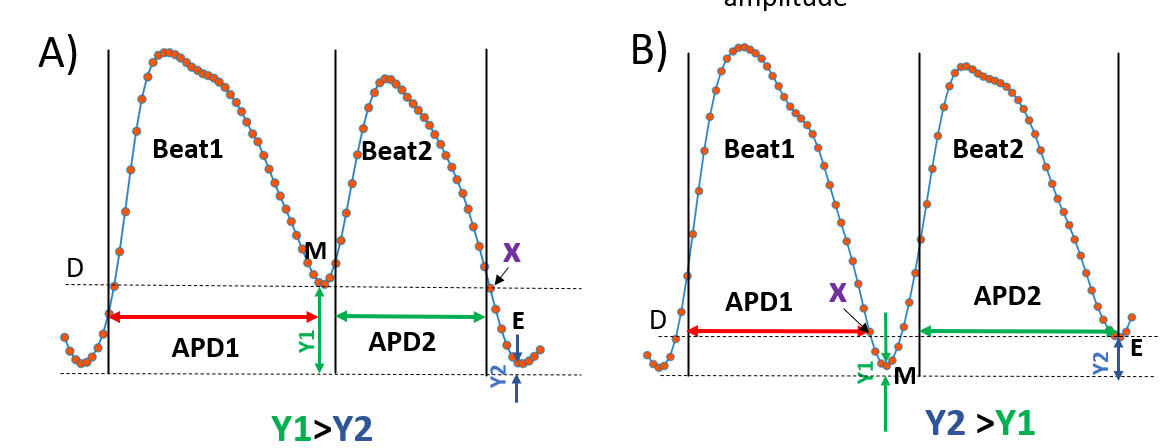

**Figure 4.** Example of “dynamic” option for AP alternan measurements.

C) User feedback AAC

A midpoint map can be erroneous when a signal is overly noisy. As shown Figure 5 near the left lateral ventricular region there are spots where M has abruptly changed to very large value than the neighbor pixels. As shown in the corresponding voltage traces the first beat (red arrow) of the selected time window exhibits biphasic plateau of the AP, which can result from noise or tissue motion. In such case, ‘User feedback AAC’ allows a user to identify the maximum M amplitude from a pop-up M-map by a mouse click. The value of maximum repolarization phase is displayed on the console. A user then uses this value in the “Fixed AAC’’ option as % repolarization, which allows AAC map at maximum repolarization.

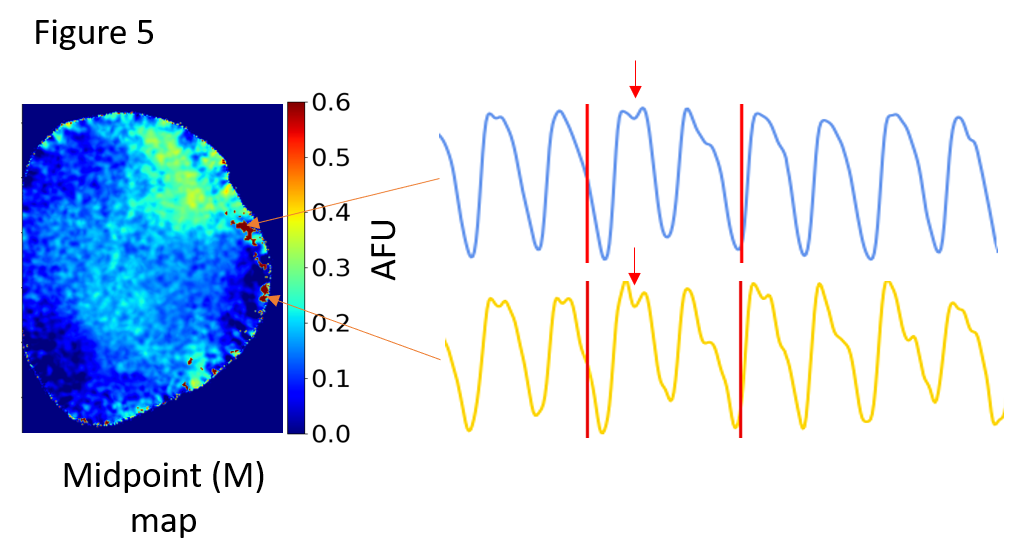

**Figure 5.** Example when midpoint map can result in errors, and the “user-defined” option for AP alternans can be used instead. AFU = arbitrary fluorescence units

**4. Extrasystolic Stimulation (S1-S2)**

To map S1-S2 paced characteristic coefficients, S1 APD is calculated from the beat that precedes the S1 beat, which is followed by the S2 beat. A user selects the time window (W1) for the first beat (Figure 6A), and APD is calculated as described in “APD and CaD mapping’’ section. Note that, it is assumed that all the S1 beats are morphologically comparable at steady state pacing. To calculate S2 APD time window W-2 is considered for the fiducial points (Figure 6A). Note that AP magnitude (normalized) reference is S1 beat in W-1. To calculate S2 APD, the algorithm first identifies the peaks (P1 and P2) of the two consecutive beat, and midpoint (M) at local minima, similar to midpoint identification in ‘Dynamic AAC and CAC’ section. The time of maximum dV/dt is identified between M and P2. Time of a specific repolarization (or reuptake, if CaT) phase (%) is identified within the time window between P2 and E.

Amplitude-based S1S2 coefficients are derived by calculating the S1 and S2 beat amplitude. Figure 6B, illustrates the measurement of the amplitudes in S1 ($\varepsilon_{{AP}_{s1}})$ and S2 ($\varepsilon_{{AP}_{s2}})$ beat of a CaT at S1S2 pacing. The algorithm finds the midpoint M as described in earlier sections.

**Figure 6.** Example of AP and CaT measurements during S1-S2 pacing.

**Image registration for ACC mapping**

Depending on the camera system used, voltage and calcium fluorescent images may need to be registered to have same coordinates of the image pixels. We described how image registration can be performed with Fiji software in the main text (for more details please see the GitHub manual). Figure 7 shows superimposed unregistered and registered voltage (red) and calcium (yellow) images.

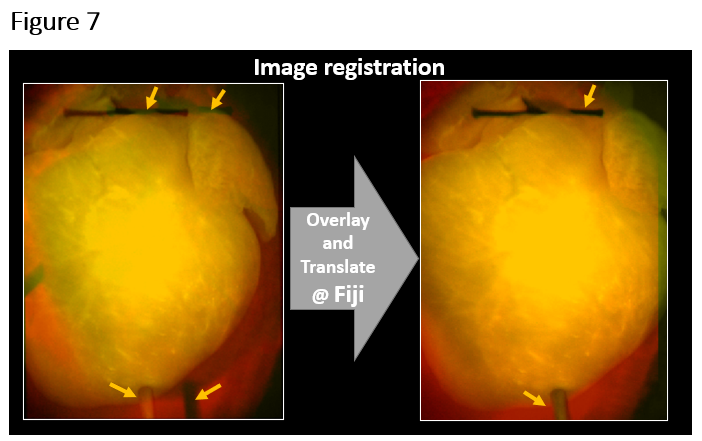

**Figure 7.** Example of image registration in Fiji.
